# The cerebellum as a moderator of negative bias of facial expression processing in depressive patients

**DOI:** 10.1101/2021.05.12.443029

**Authors:** Anna Nakamura, Yukihito Yomogida, Miho Ota, Junko Matsuo, Ikki Ishida, Shinsuke Hidese, Hiroshi Kunugi

## Abstract

**Background:** Negative bias—a mood-congruent bias in emotion processing—is an important aspect of major depressive disorder (MDD), and such a bias in facial expression recognition has a significant effect on patients’ social lives. Neuroscience research shows abnormal activity in emotion-processing systems regarding facial expressions in MDD. However, the neural basis of negative bias in facial expression processing has not been explored directly.

**Methods:** Sixteen patients with MDD and twenty-three healthy controls (HC) who underwent an fMRI scan during an explicit facial emotion task with happy to sad faces were selected. We identified brain areas in which the MDD and HC groups showed different correlations between the behavioral negative bias scores and functional activities.

**Results:** Behavioral data confirmed the existence of a higher negative bias in the MDD group. Regarding the relationship with neural activity, higher activity of happy faces in the posterior cerebellum was related to a higher negative bias in the MDD group, but lower negative bias in the HC group.

**Limitations:** The sample size was small, and the possible effects of medication were not controlled for in this study.

**Conclusions:** We confirmed a negative bias in the recognition of facial expressions in patients with MDD. fMRI data suggest the cerebellum as a moderator of facial emotion processing, which biases the recognition of facial expressions toward their own mood.

## 1. Introduction

Social functioning relies on recognizing the emotions of people, but this functioning is impaired in those with major depressive disorder (MDD) (Joiner, 2002). Cumulative studies confirm a mood-congruent emotion-processing bias, that is, a “negative bias” to be a core aspect of MDD (e.g., Everaert et al., 2012). The difficulties in understanding nonverbal cues may undermine the satisfaction of interpersonal communication (Carton et al., 1999), leading to negative social ties and the development of clinical depression (Finch and Zautra, 1992).

Facial expressions play a crucial role in emotion recognition (Ekman, 1999; Keltner et al., 2003). A number of studies support the existence of a negative bias in facial expression recognition in patients with MDD (reviewed in Bourke et al., 2010). For example, the MDD group tended to label neutral faces as sad in Leppänen et al.’s (2004) study, and the increase in depressive mood in MDD patients resulted in higher precision of sad faces compared with healthy volunteers (Gollan et al., 2010). Joormann and Gotlib (2006) found that patients with MDD needed more intensive expression to recognize happy emotions, while they recognized sad emotions from slighter expressions, compared with healthy controls.

The neural response to facial expressions has been intensively studied to understand the background of the emotional and cognitive symptoms in depression; however, to our knowledge, none of the studies have directly examined the brain function related to the negative bias. Studies using noninvasive neuroimaging methods, such as functional magnetic resonance imaging (fMRI), show abnormal activity in the emotion-processing network in depression, especially in the limbic system (amygdala, hippocampus, insula, thalamus, and striatum) (Fitzgerald et al., 2008). Groenewold et al. (2013) reports the limbic region’s hyperactivity in response to negative faces, its hypoactivity for positive faces. Abnormalities in frontal and temporal activity, as well as connectivity between the frontal area and amygdala, have also been reported (Stuhrmann et al., 2011). To assess the neural response to facial expressions, scholars have typically employed implicit (e.g., asking the gender of the image of a model showing facial expression), subliminal (e.g., showing facial expressions for 25–40 ms followed by a neutral face), or matching (e.g., asking to match sad faces with a fearful distracter) tasks. Unfortunately, these task paradigms make it difficult to directly examine the neural basis of the negative bias in facial expression recognition, as they do not require the participants to explicitly judge the emotion in the expressions. Although Almeida et al. (2010) and Lee et al. (2008) employed explicit tasks (e.g., asking whether the face is emotional or not), they did not examine the relationship between brain activity and negative bias. While Ito et al. (2017) employed explicit tasks and reported that brain responses to facial stimuli in the anterior cingulate cortex were associated with negative bias, their participants were limited to healthy individuals. Therefore, the neuropathology of negative bias in MDD remains elusive.

In the present study, our primary goal was to reveal the neural basis of the negative bias in MDD. While undergoing fMRI, both patients with MDD and healthy controls completed an explicit facial task that measures the degree of negative bias. We tried to identify brain regions whose activity was associated with the degree of negative bias, and a group comparison was conducted to clarify MDD-specific neuropathology.

## 2. Methods

### 2.1. Participants

This study was performed in accordance with the Declaration of Helsinki (World Medical Association, 2013). The study protocol was approved by the Research Ethics Committee of the National Center of Neurology and Psychiatry (Tokyo, Japan), and all participants provided written informed consent for their participation. Eighteen Japanese patients with MDD and twenty-four healthy volunteers (HC) were recruited with the following conditions: right-handed, aged between 20 and 70 years, and no contraindication to MRI scanning (e.g., metallic implants or claustrophobia). Three patients were excluded owing to excessive head motion during the fMRI scan.

Consensus diagnoses were determined according to the criteria laid out in the Diagnostic and Statistical Manual of Mental Disorders, 5th edition (DSM-5; American Psychiatric Association, 2013), based on the information from the Mini International Neuropsychiatric Interview (Sheehan et al., 1998; translated by Otsubo et al., 2005), additional unstructured interviews, and medical records, if available. The healthy controls had no history of contact with any psychiatric services. Participants were excluded if they had a medical history of central nervous system diseases, severe head injury, substance abuse, or mental retardation. Eventually, the data of 16 MDD and 23 HC participants were included in the analysis (Table 1).

**Table 1.**
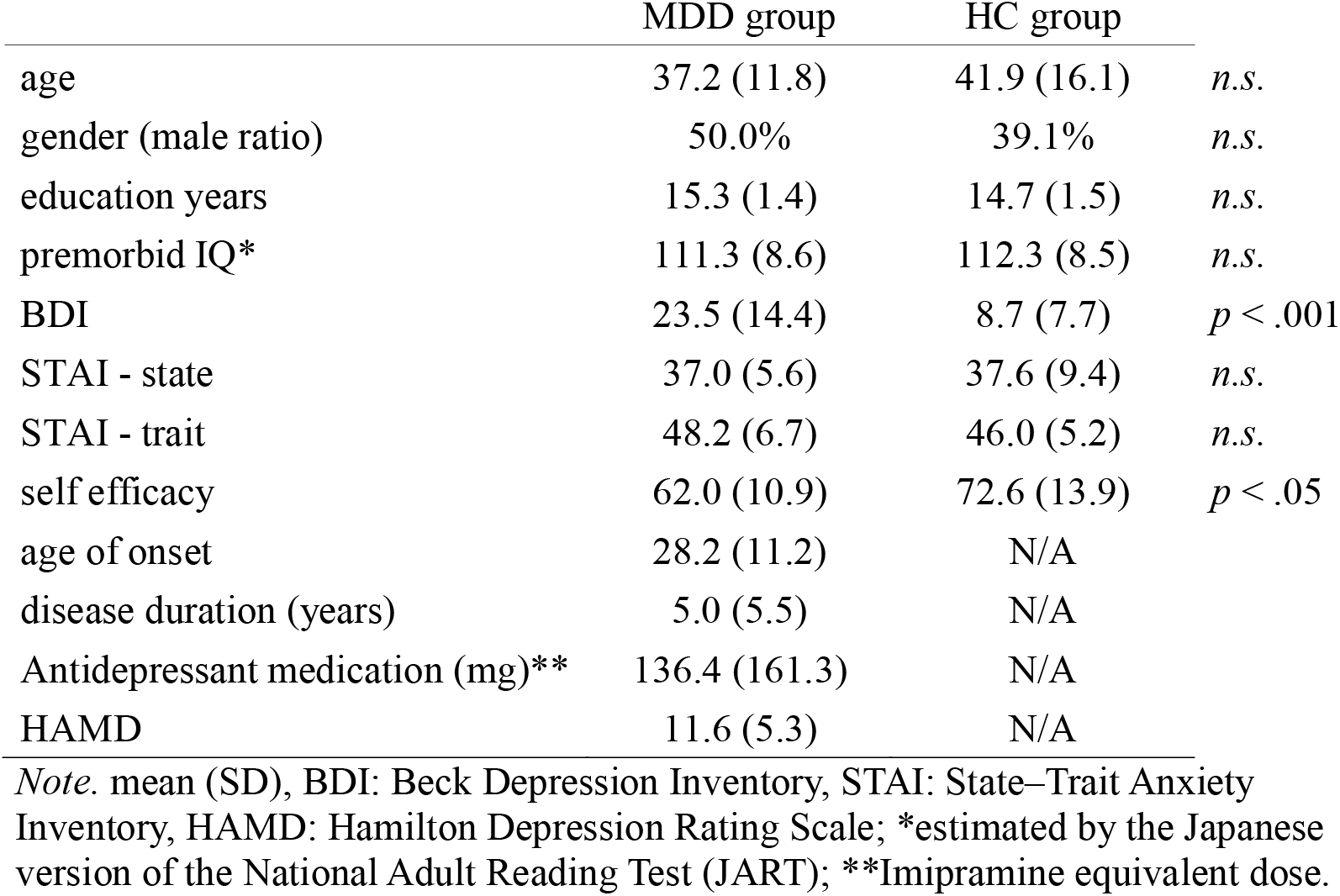
Demographic data of MDD and HC groups

MDD patients responded to 21 items on the GRID–Hamilton Depression Rating Scale (HAM-D) Williams et al., 2008; Japanese version translated by Nakane, 2004) to evaluate the severity of the symptoms. Their daily doses of antidepressants were converted to imipramine equivalents using published guidelines (Inada and Inagaki, 2015). All participants completed a self-report questionnaire including the second version of the Beck Depression Inventory (BDI-II) Beck et al., 1996; Japanese version translated by Kojima et al., 2002), the Japanese version of the generalized self-efficacy scale (Narita et al, 1995), and the State–Trait Anxiety Inventory (STAI) (Fountoulakis et al., 2006; translated by Iwata et al., 1998). The premorbid IQ scores of all participants were estimated using the Japanese version of the National Adult Reading Test (Matsuoka et al., 2006).

### 2.2. Measurements: Facial Task

#### 2.2.1. Stimuli

The Taiwanese Facial Expression Image Database (TFEID), produced by the National Yang-Ming University, was used to generate a facial image set (Chen and Yen, 2007). We chose this database that contains the greatest number of Asian facial images because facial expression processing is suppressed when responding to faces of a different race from ourselves (Malpass and Kravitz, 1969). From the TFEID set, we selected the facial images of 20 models for the categories of “happiness,” “neutral,” and “sadness.” To manipulate the intensity of the expressions, we combined neutral images with sad or happy images to generate the 25%, 50%, and 75% conditions using a standardized procedure to morph facial expressions (Young et al., 1997). Norrkross MorphX (Norrkross Software) was used for this process. Consequently, nine types of images (happy–100%, happy–75%, happy–50%, happy–25%, neutral, sad–25%, sad–50%, sad–75%, and sad–100%) were produced for each model.

#### 2.2.2. Task Procedure

The facial task was prepared and presented using MATLAB (version R2018a, Mathworks) with Psychophysics Toolbox extensions (version 3.0.14) (Brainard, 1997; Kleiner et al., 2007; Pelli, 1997). In each trial, participants were instructed to stare at a fixation point for 3–5 s; subsequently, a facial expression image was shown for 500 ms each (*see* Figure 1).

**Figure 1.**
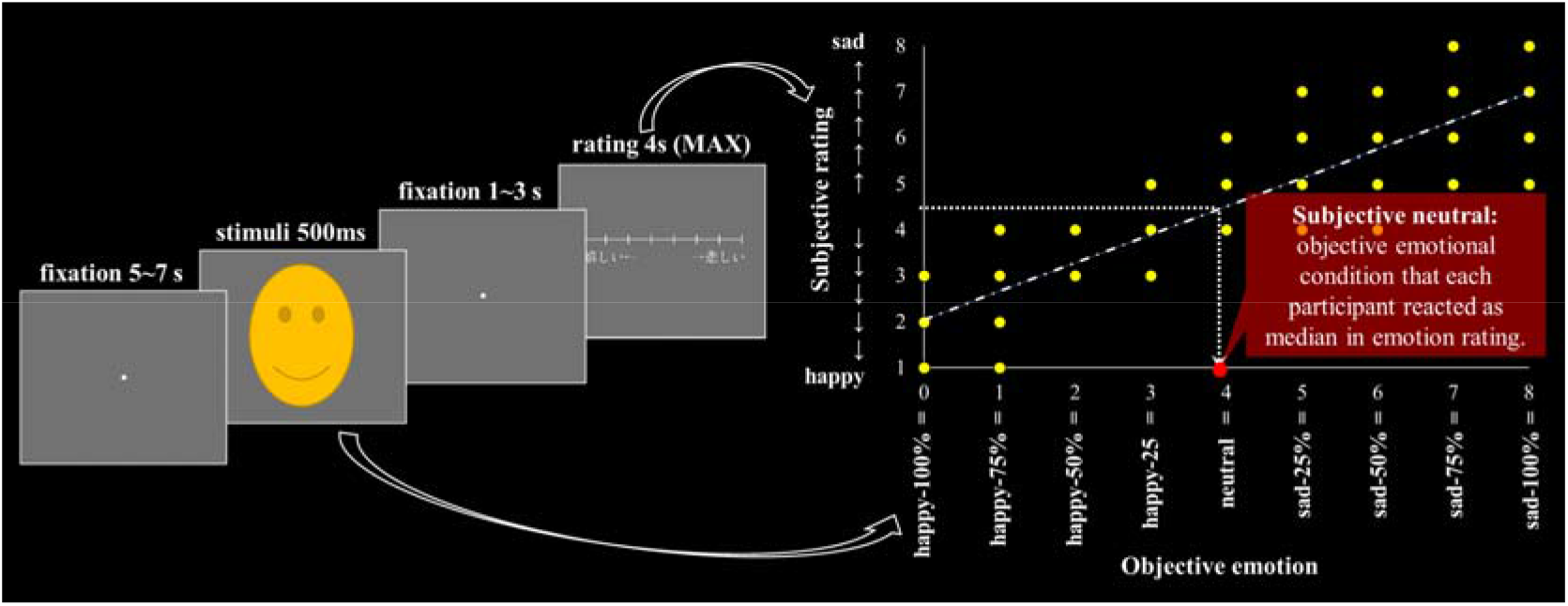
Task procedure of facial task and the relationship between variables. *Note*. A facial stimulus was chosen randomly from one of the nine emotional conditions: happy, 100%; happy, 75%; happy, 50%; happy, 25%; neutral, sad, 25%; sad, 50%; and sad, 75%, and 100%. In the rating phase, the participants were asked to rate the face with an eight-point scale, ranging from happy (嬉しい) to sad (悲しい), using the prescribed keys. The rating phase ended 1 s after the key was pressed when the participant answered, or it automatically moved to the next trial after 4 s of waiting, with no key being pressed. The participants completed 180 trials in total and were divided into two blocks. The illustration of the face is different from the actual stimuli.

After seeing the image, the participants were required to rate the emotion of the expression using an eight-point scale on the response bar, from “happy” to “sad” (1 = very happy and 8 = very sad). The participants were required to indicate their answers by pressing the prescribed keys. The task automatically moved to the next trial either 1 s after the participant pressed the key or 4 s after the response bar was shown, if the participant had not decided any answer. Each participant performed 180 trials, one trial per image: nine conditions (happy–100%, happy–75%, happy–50%, happy–25%, neutral, sad–25%, sad–50%, sad–75%, and sad–100%) × 20 models. Trials were divided into two blocks, and one scan session was allotted for each block, including a rest in the middle. The right and left sides of the response bar were flipped between blocks for each participant to counterbalance the association of the rating scale and the participants’ hand/finger. Ninety trials were presented in random order in each block, and the order of the two blocks was counterbalanced between the subjects. The participants performed 10 practice trials, 5 trials both outside and inside the MRI scanner, before the actual task with fMRI scanning. The participants were reimbursed for their time and any other expenses related to the experiment.

#### 2.2.3. Converting Variable

To measure the negative bias in facial expression processing, we calculated the “subjective neutral” score for each participant. Subjective neutral reflects the objective expression that the participant subjectively felt as neutral (Figure 1: right). Objectively happier “subjective neutral” means that the individual had a negative bias in rating facial expressions, and vice versa. We fit a linear regression model *Y* = *aX* + *b* to the data for which the *X* value is the objective emotion score (0 = happy–100%, 4 = neutral, and 8 = sad–100%) and the *Y* value is the subjective rating score (1 = very happy and 8 = very sad). After estimating the linear regression, we substituted the median of the range of subjective rating (4.5) for *Y*, which corresponds to the point where the subject reacted as neutral, to the formula. We called the *X* gained from that calculation “subjective neutral.” Although Ito et al. (2017) used ordered logistic regression to fit their data for the rating of morphed facial expressions, we applied linear regression as there are differences in the rating (two-alternative forced-choice or eight-point scale) and the emotions used (morphing happy with sad or with neutral). The R^2^ of linear regression and logistic regression of each participant’s data were not significantly different (*t*(76) = .36, *p* = .72).

### 2.3. fMRI Image Acquisition

Functional imaging was conducted using a 3-Tesla MR system (Trio, Siemens, Erlangen, Germany) to acquire gradient T2*-weighted echo-planar images (EPI) with blood oxygenation level-dependent contrasts. Forty-two contiguously interleaved transverse EPI-image slices were acquired in each volume (slice thickness, 3 mm; no gap; repetition time, 3,000 ms; echo time, 25 ms; flip angle, 83°; field of view, 192 mm ^2^; matrix, 64 × 64). We discarded the first three volumes to compensate for T1 saturation effects. A high-resolution anatomical T1-weighted image was acquired for each subject.

### 2.4. Statistical Process and Analysis

The fMRI data were preprocessed and analyzed using SPM12 software (Welcome Center for Human Neuroimaging, London, UK) implemented in MATLAB R2018a. MarsBaR (0.44) was used to calculate the contrast estimate of the specific area as a toolbox on SPM12. SPSS Statistics 24.0 (IBM) and Microsoft Excel were used for the clinical and behavioral analyses.

Regarding behavioral data, we conducted a two-sample t-test with the “subjective neutral” measure to see the difference in negative bias between the MDD and HC groups. Following the literature, we expected that the MDD group would show a higher negative bias compared with the HC group.

Regarding the fMRI data, the following preprocessing measures were performed: correcting for head motion, adjusting acquisition timing across slices, spatial normalization to the standard Montreal Neurological Institute (MNI) template, and smoothing using a Gaussian kernel with a full-width at half the maximum of 8 mm. After preprocessing, a conventional two-level approach was adopted using the SPM12. Regressors were generated for each condition (i.e., happy, neutral, sad) by convolving a canonical hemodynamic response function provided by SPM12 with the onset time of face presentation (500-ms duration). Key pressing, visual stimuli other than faces (response bars), and head movements computed through the realignment process were also included in the model as regressors of no interest. To explore the brain activation pattern associated with the negativity bias and its group differences, we first created contrasts [happy > neutral] and [sad > neutral] to extract brain activity specific to processing each emotional expression. For each emotional category, these activations were entered into group-level multiple regression analyses, which related them to the degree of negativity bias (i.e., subjective neutral score) of the MDD and HC groups. Brain regions showing group differences were assessed with the statistical threshold set at an uncorrected p < 0.001 at the voxel level and at p < 0.05, family wise error (FWE) corrected at the cluster level, assuming the whole brain as the search volume.

## 3. Results

There were no significant differences in age, sex ratio, years of education, or premorbid IQ scores between the two groups (*see* Table 1). The MDD group had significantly higher BDI-II scores (*t*(37) = 4.16, *p* < .0005, *d* = 1.35) and significantly lower self-efficacy scores (*t*(37) = 2.54, *p* < .05, *d* = 0.83) compared with the HC group. The STAI scores were not significantly different between the two groups (state: *t*(37) = 0.23, *p* = .81, *d* = .08 ; trait: *t*(37) = 1.17, *p* = .24, *d* = .38). The mean HAM-D score of the MDD group was 11.6 (SD = 5.3), indicating mild depression.

Behaviorally, a two-sample t-test revealed that the MDD group recognized happier faces as subjectively neutral compared with the HC group (*t(*37) = 2.25, *p* = .03, *d* = .73), which confirmed the presence of negative bias.

The brain area that correlated with the negative bias was then examined. The whole-brain analysis showed that the MDD and HC groups showed a different pattern of correlation between subjective neutral and brain activity in response to happy faces in left cerebellum lobule VI (peak MNI coordinates −22, −74, −22; *p* < .05, FEW whole-brain corrected; Figure 2). While the stronger response to happy faces in this region was related to lower negative bias (higher positive bias) in the HC group (*r* = .65), the same tendency of cerebellar activity corresponded to a higher negative bias in the MDD group (*r* = −.59). The activity of sad *versus* neutral faces did not correlate with the negative bias in any brain region.

**Figure 2.**
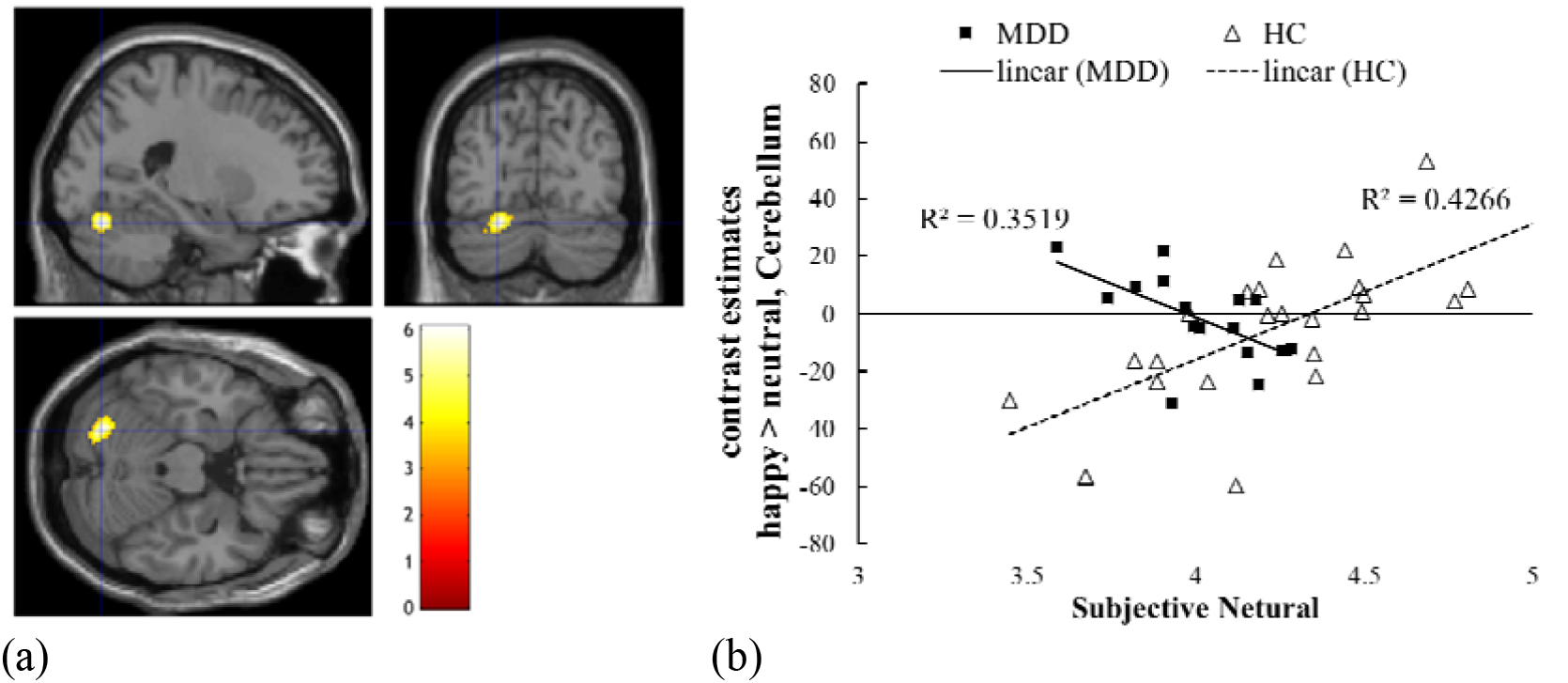
Interaction of correlation between activation in cerebellum lobule VI at happy > neutral and the behavioral negative bias in MDD and HC groups. *Note*. (a) GLM with covariate of interest FWE p = .05, (b) correlation of subjective neutral and contrast estimate of 8 mm hemisphere around the peak coordinate shown in (a). * Low subjective neutrality corresponds to a stronger negative bias.

## 4. Discussion and Conclusion

The current study aimed to reveal the neural basis of negative bias in patients with MDD. As the literature suggests, our MDD patients showed a mood-congruent bias in facial expression processing. They recognized facial expressions in a negatively biased way, reflecting their negative mood. Our findings suggest that the posterior cerebellum plays an important role in this process. Our notion is supported by the following previous findings that the cerebellum plays an important role in higher cognitive functions.

The cerebellum has traditionally been conceived as a modulator of motor functions, while recent evidence shows its engagement in cognitive and emotional processing, including recognition of facial expression (Van Overwalle et al., 2020). In summary, Schmahmann and Sherman (1998) first pointed out that patients with a lesion in the posterior cerebellum show deficits not only in their motor function but also in cognitive and emotional functions. A meta-analysis of functional neuroimaging studies confirmed that the posterior cerebellum is engaged in emotion processing (Stoodley and Schmahmann, 2009). This emotional processing in the cerebellum seems multimodal. For example, Adamaszek et al. (2014) reported that patients with cerebellar lesions had lower precision in recognizing emotions from both facial expressions and prosody. Regarding facial emotion processing in the cerebellum, a recent study provides causal evidence that applying transcranial magnetic stimulation to the left cerebellum leads to deficits in both explicit and implicit facial expression tasks (Ferrari et al., 2018). These findings support the idea that the cerebellum, especially the left posterior area found in this study, is involved in facial emotion processing.

In addition to emotion processing, a part of the posterior cerebellum (lobule VI and Crus I) was found to be related to self-referencing (Van Overwalle et al., 2014). Self-referencing is commonly measured with a task that asks participants to judge whether presented trait words describe themselves (e.g., “In general, how brave are you?”), reflecting the ability to understand the mental state of a person. Van Overwalle et al. (2020) further suggest that the posterior cerebellum is involved in the emotional aspect of self-experiences, which is measured by questioning their mood states (e.g., “How are you feeling?”). Clausi et al. (2019) reported that patients with MDD and cerebellar lesions have difficulty recognizing their own negative mood. These studies indicate that referencing self-mood is served by the cerebellum.

Taken together, previous findings indicate that the cerebellum, especially its posterior part, plays a key role in processing both facial expression and mood. This suggests that the posterior cerebellum might play a role in biasing the recognition of facial emotions toward their own mood. This assumption can explain why MDD participants with higher activity in the posterior cerebellum showed a higher degree of negative bias in the present study.

However, our assumption predicts an opposite pattern in the HC group. As participants in the HC group would generally have a non-negative (i.e., neutral to positive) mood, the higher activity in the cerebellum should bring about more positively biased facial recognition, which is manifested as a low degree of negative bias in our paradigm. This is indeed what we found. Our assumption that the cerebellum plays a role in modulating emotional processing is further supported by previous findings that this region engages in emotional regulation (Parvizi et al., 2001; Schutter and van Honk, 2009; Steward et al., 2016). Ito et al. (2017) reported that the brain response to facial stimuli in the anterior cingulate cortex is associated with the degree of negative bias observed in healthy participants. The difference between their findings and ours can be attributed to the difference in the focus of the analysis. In the present study, we focused on the difference in brain activity associated with a negative bias between HC and MCC groups in order to clarify neuropathology specific to MDD. In this sense, our findings do not necessarily conflict with the findings of Ito et al.. Our novel finding regarding cerebellar involvement in negative bias may contribute to expanding the neural account of this important phenomenon.

It has long been pointed out that the cerebellum of MDD patients has abnormal structure and function (Adamaszek et al., 2017, as a review), and abnormalities in the cerebellum have been deemed to have a causal role in depression (D’Angelo and Casali, 2013; Schmahmann, 2004). A meta-analysis revealed that patients with MDD show altered cerebellar responses to emotional stimuli (Fitzgerald et al., 2008) and abnormal connectivity in the large-scale cerebro-cerebellar pathways (Liston et al., 2014; Wang et al., 2017; Zang et al., 2012). These cerebellar abnormalities could have partly reflected the neuropathology related to the negative bias that we found here. An important avenue for future research on negative bias will be to explore its relation to the connectivity between the cerebellum and the limbic/frontal system, where studies suggest that it plays a pivotal role in mood disorders (Dean and Keshavan, 2017; Fitzgerald et al., 2008; Price and Drevets, 2010).

The present study was limited by its sample size, which may have caused false-negative errors. As most of the MDD participants were under antidepressant medication, mainly SSRIs and SNRIs, it is possible that medication may have effects on the results of this study. Further studies with a larger sample size while controlling the medication will be needed.

Despite these limitations, our study showed a distinct relationship between cerebellar activity and negative bias at the behavioral level. As mood-congruent bias is found in many aspects of cognition, such as memory, attention, and interpretation. Further, investigation is expected to determine the validity of our hypothesis that the negative bias derives from self-referencing others’ emotions with their own mood in the cerebellum. The findings of this study have important implications for future research on the pathology of depression and have the potential to be applied to the assessment of and intervention in MDD patients in clinical practice.

## Acknowledgements

We would like to thank Editage (www.editage.com) for English language editing.

